# Intranasal oxytocin drives coordinated social approach

**DOI:** 10.1101/2020.11.20.390245

**Authors:** Patrick K. Monari, Nathaniel S. Rieger, Juliette Schefelker, Catherine A. Marler

**Affiliations:** Department of Psychology, University of Wisconsin-Madison, Madison, WI, USA; Department of Psychology and Neuroscience, Boston College, Chestnut Hill, MA, USA

**Keywords:** Oxytocin, intranasal oxytocin, pair bonding, coordination, approach, ultrasonic vocalizations, *Peromyscus californicus*, California mouse

## Abstract

Coordinated responses to challenge are essential to survival for bonded monogamous animals and may depend on behavioral compatibility. Oxytocin (OT) context-dependently regulates social affiliation and vocal communication, but its role in pair members’ decision to jointly respond to challenge is unclear. To test for OT effects, California mouse females received an intranasal dose of OT (IN-OT) or saline after bonding with males either matched or in their approach response to an aggressive vocal challenge. Pair mates were re-tested jointly for approach response, time spent together, and vocalizations. Females and males converged in their approach after pairing, but mismatched pairs with females given a single dose of IN-OT displayed a greater convergence that resulted from behavioral changes by both pair members. Unpaired females given IN-OT did not change their approach, indicating a social partner was necessary for effects to emerge. Moreover, IN-OT increased time spent approaching together, suggesting behavioral coordination beyond a further increase in bonding. This OT-induced increase in joint approach was associated with a decrease in the proportion of sustained vocalizations, a type of vocalization that can be associated with intra-pair conflict. Our results expand OT’s effects on behavioral coordination and underscore the importance of emergent social context.

## Introduction

Coordinated behavior is essential to compatibility and survival for species that are biparental and develop mating pair bonds. Examples across vertebrate taxa illustrate environmental and social challenges that require joint strategies by bonded partners [1–3], and for monogamous and territorial species one such challenge is the introduction of an unfamiliar conspecific. Pairs with an established bond must appropriately recognize and confront outgroup individuals, as their presence threatens territorial infringement [4], resource theft [5], and infanticide [6], all of which can hamper pair fitness [7–9]. Some species, such as cichlid fish [10] and Kirk’s dik-dik [11] respond with rigid, sex-specific behavioral roles such that one sex invariably defends territories while the other cares for offspring. Other pair bond-forming species, however, display more flexible behavior between sexes. For example, prairie vole [12,13] and California mouse [14–16] pair mates closely overlap in their behavior, and both sexes can complete any necessary task besides nursing. Compatible responses between pair-bonded individuals to an extrapair challenge are therefore uniquely important to coordination in these species.

The neuropeptide oxytocin (OT) context-dependently regulates social affiliation [17] and vocal communication [18], and by extension may impact response compatibility, but its role in pair members’ decision to jointly react to challenge is unclear. OT signaling facilitates prosocial behaviors in some cases, while it promotes social vigilance [19], social anxiety [20], and agonistic behaviors [21,22] in others. The effects of OT on investigation and aggression are therefore complex and rely on social context [23] and may modulate behavior and emotion by enhancing an individual’s attunement to its social landscape [24–26]. For example, in marmosets, intranasal OT (IN-OT) increased approach towards an unfamiliar conspecific when a mate was also present but decreased approach when the mate was absent [27]. In humans, IN-OT reduced xenophobic aggression when paired with injunctive prosocial norms of peers, but not when administered without them [28]. Taken together, this evidence supports the notion that OT signaling influences how social affiliations moderate responses to the outgroup.

An outstanding question is whether, and under what circumstances, OT facilitates coordination of approach toward an unfamiliar conspecific. California mice (*Peromyscus californicus*) are uniquely positioned to answer this question as biparental, highly territorial mammals that flexibly cooperate within the pair bond to accomplish goals such as intruder investigation [16,29]. In the wild and the laboratory, adults form lifelong female-male dyads and are well documented to be highly aggressive toward other same-sex or opposite-sex individuals [14,16,30–36], restricting the ecological relevance of cooperative behavior to mating pairs and possibly family units. Paired California mice can show both synchronous or divided defense in response to intrusion, with varying degrees of individual and joint approach by pair members, suggesting variability in pair coordination and division of labor [37]. Flexible application of coordinated strategies may depend on the type and degree of challenge; in some cases, joint action may help to efficiently remove a threat, while in other cases it might be advantageous to divide location and simultaneously defend resources or pups while addressing an intruder. We recently found that, prior to pup birth and following pair-bonding, individuals of both sexes adjusted to match their partner’s level of approach in a compatibility-dependent manner when exposed to a low level of challenge; paired individuals that were mismatched (one individual higher, one lower) in their approach before pairing were more likely to change their behavior to become similar after pairing, while matched pairs (similar levels of approach between individuals) changed little. The change in behavior for mismatched pairs was associated with changes to ultrasonic vocalization (USV) production profiles [38, in review]. We speculate that vocal communication plays a role in the decision to adopt one coordination type over the other, as paired California mice display a rich variety of USVs that have been linked to complex social behavior [39,40]. However, the neuroendocrine mechanisms underlying this coordination-induced change to social approach have yet to be studied.

OT can context-dependently regulate approach and avoidance of a novel conspecific in female California mice [41,42]. Here we used female IN-OT administration and pre-pairing/post-pairing simulated intrusions, via playbacks of aggressive vocalizations, to show that OT is critical to mediating pair-bonding-induced changes to social approach. Two hypotheses emerge for how OT may modulate pair coordination in response to challenge: the convergence hypothesis predicts that OT will increase the similarity between pairs mismatched in their degree of social approach, increasing the amount of time pair mates spend jointly addressing an unfamiliar intruder [43,44]. Conversely, the divergence hypothesis predicts that OT can increase the dissimilarity between matched pairs, increasing the amount of time pair mates spend apart, thereby dividing labor [37].

## Methods

### Animals

48 adult males and 48 adult females (age 3-6 months) were housed in standard cages (48 × 27 × 16 cm) lined with aspen bedding and a nestlet with Purina 5015™ mouse chow and water available *ad libitum*, 2-4 same-sex individuals per cage. All tests occurred between 1–3 hrs after the onset of the dark cycle in dim red light in housing maintained at 20–23° C on a 14:10 h light:dark cycle (lights on at 16:00 central standard time). Males and females were tested for response to bark playbacks and were selectively paired and housed in new cages. In a separate experiment, 20 females were left unpaired and housed with their original cage mates.

### Ethical statement

Animals were maintained according to the National Institute of Health *Guide for the Care and Use of Laboratory Animals*. Procedures were approved by the University of Wisconsin-Madison College of Letters and Sciences Institutional Animal Care and Use Committee (Protocol L005447). No animals were injured by any of the behavioral manipulations or assays.

### Apparatus

Testing occurred in aspen bedding-lined Plexiglas cages (90 × 30 × 30 cm) equally divided into three chambers (each 30 × 30 × 30 cm) with centrally located openings (11.5 × 11.5 cm) between chambers to allow for free movement. Speakers (Vifa Dynamic Ultrasound, 1-120 kHz range, Avisoft Bioacoustics, Berlin, Germany) were placed at each end of the apparatus against a closed mesh gate.

### Playback tracks

We investigated California mouse approach behavior towards playbacks of loud, aversive bark calls, adapted from [38, in review]. In a separate cohort used exclusively to produce the playback stimuli, individual male and female mice were placed in a single-chambered plexiglass apparatus (50 × 30 × 30) under normal food and water conditions for 24 hrs, a length of time demonstrated to be sufficient for the formation of residency behavior, and in which the arena becomes the individual’s territory [36]. After 24 hrs, residents were introduced to a same-sex intruder for an 8-min aggressive encounter period similar to previous studies [32,33,36,45,46]. Each intruder was only used for a single encounter, and otherwise had no previous aggression testing experience. During the encounter we recorded defensive-aggressive barks using an ultrasonic microphone (Emkay/Knowles FG series, detection range: 10-120 kHz) with a 250 kHz sampling rate and 16-bit resolution, placed 30cm above the bottom of the apparatus. Spectrograms were produced using a 512 fast Fourier transform in Avisoft SASlab pro (Avisoft Bioacoustics, Berlin, Germany) in order to identify barks. Barks appear as short, high-amplitude calls with an upside down U shape that begins and ends in the audible range for humans [16,47] (Fig. 1A). We created playback tracks using these spectrograms by selecting only bark calls. Calls could not be distinguished between the resident and the intruder during the encounter, therefore both resident and intruder barks were used to construct playback tracks. Playback tracks were 2 mins in duration and contained 120 ± 5 bark calls. Output gain/volume was maintained across playback tracks. The ambient noise track control was a 2 min recording of the quiet testing room with all lights off and no mice present. We used 8 unique tracks from 8 different sets of individuals and assigned tracks to individuals randomly with each track used between 20-23 times [38, in review], ensuring that no individual heard the same track more than once over the course of the two tests (to avoid habituation and maintain consistency).

**Figure 1.**
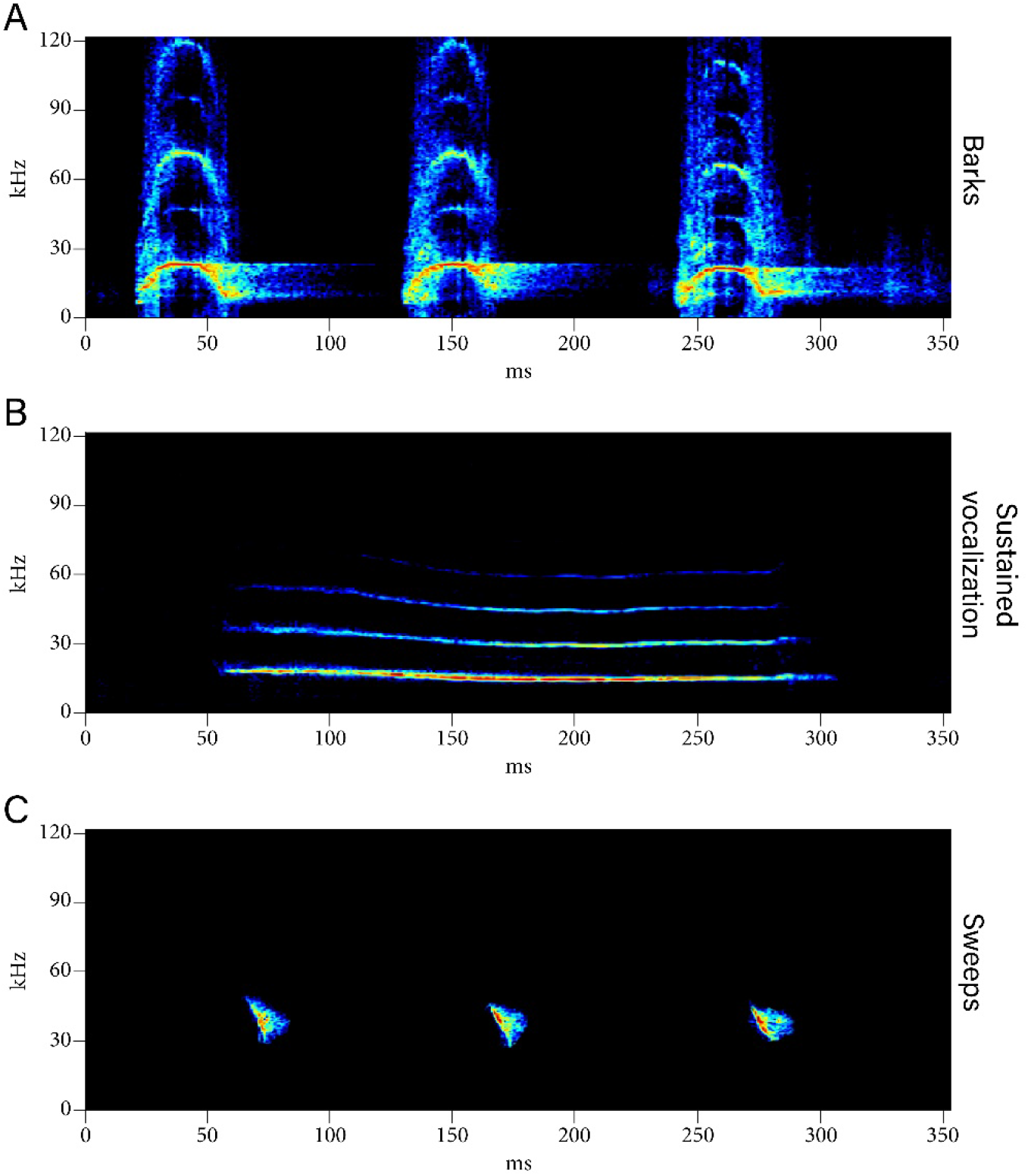
California mouse vocalizations. Examples of **A**. Barks, **B**. A sustained vocalization (SV), and **C**. Sweeps.

### Pre-pairing playback approach test

Mice were first tested for response to bark playbacks as nonbonded, sexually naïve individuals (Fig. 2A, left). Mice were placed in the testing apparatus for 5-10 mins to habituate to all three chambers. One male and two females did not enter all three chambers by 10-min and so were removed from the experiment (resulting in 47 males and 46 females). Two-min playback tracks were played from speakers at opposite ends of the apparatus behind wire mesh, with one speaker playing a bark track and the other an ambient noise track concurrently. The bark track and ambient noise track speaker locations were randomized across trials. Video and audio recordings were made of their behavior. We recorded time spent in the chamber closest to the bark speaker (“approach chamber”) as an approach score, and aggregate time spent in the chamber closest to the ambient noise speaker (“avoid chamber”) as an avoidance score.

**Figure 2.**
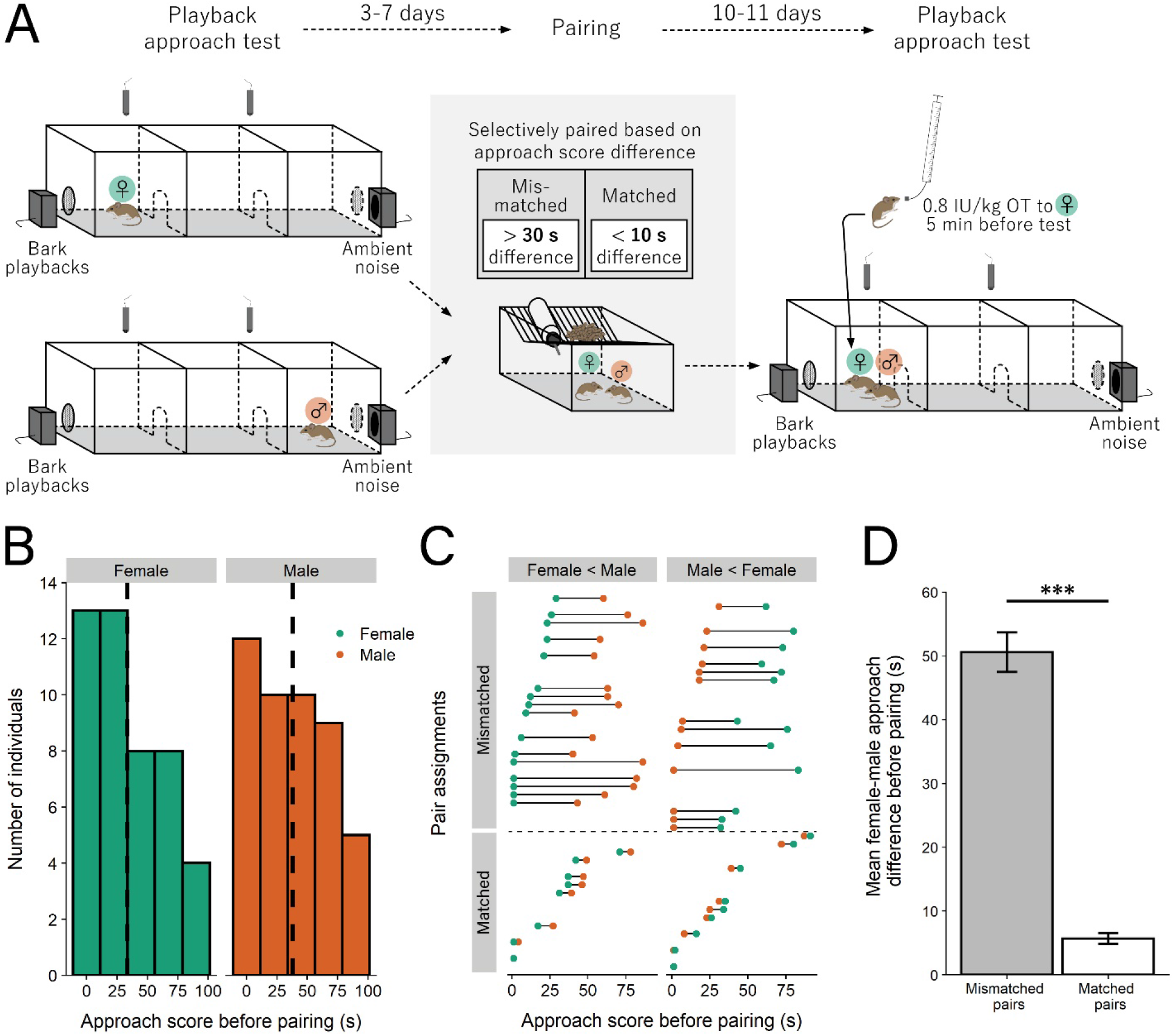
Selective pairing to form mismatched and matched pairs based on initial social approach score. **A**. Experimental design, adapted from [38, in review]. Approach is the amount of time spent in the chamber closet to the bark playback speaker. **B**. Distribution of approach scores for females and males before pairing; dotted line is mean approach. **C**. Stratified randomized pair assignments; mismatched pairs had ≥ 30 s difference in approach score between mates while matched pairs had ≤ 10 s difference. **D**. The female-male approch difference for mismatched pairs was significantly greater than that for matched pairs (*** = p < 0.001). Error bars are ± s.e.m.

### Behavioral type and pairing

3-7 days following the pre-pairing test, 46 females were selectively paired with 46 males in a stratified randomized design based on approach scores. Mice were paired such that they were “matched” in their approach scores (i.e., their approach scores were ≤ 10 s different from each other, N = 17) or they were “mismatched” (i.e., their approach scores were ≥ 30 s different from each other, N = 29). There were no pairs formed that had an approach score difference between 10 and 30 s. In a follow-up experiment, 20 female mice were left unpaired following the first playback approach test.

### Post-pairing playback approach test

All mice underwent a second playback test to determine if pairing alters responses to bark calls. Pairs were retested 10-11 days after pairing (13-17 days after the pre-pairing test). At 7 days post-pairing, pairs exhibit hallmarks of pair-bonding [48], including increased affiliation and decreased aggression, indicating that 10-11 days is sufficient for pair-bond formation. The playback procedure was the same as in the pre-pairing test except that paired mice were tested together as a pair (Fig. 2A, right). Both mice were placed into the central chamber and required to enter all three chambers prior to testing. Behavior scoring was the same as in the first test and, additionally, pairs were scored for time spent together in the same chamber. Females generally give birth within 35 days of pairing when housed in cages of similar size to those used in this experiment [37], so were expected to be pregnant at the time of the post-pairing test (gestation 31-32 days, [49]). The procedure for unpaired mice was the same as the pre-pairing test. Two unpaired females were removed as they did not enter all three chambers by 10 mins (resulting in 18 females).

### IN-OT dose and application

We administered a 0.8 IU/kg dose of IN-OT (Bachem, Torrance, CA, Prod #: 4016373). This dose induces changes in female behavior in rodents [50] and approximates a weight-adjusted dose used in human studies [51]. Intranasal administration is an established, non-invasive route of delivery for OT [52], and intranasally-administered OT enters the brain in house mice (*Mus musculus*) at behaviorally-relevant concentrations [53]. In California mice, IN-OT produces behavioral effects similar to centrally-administered OT, suggesting that it reaches the brain in this species [41,54]. Following pairing, mice were randomly assigned to either saline (mismatched pairs: N = 15; matched pairs: N = 8) or IN-OT treatment (mismatched pairs: N = 14; matched pairs: N = 9) groups. Unpaired females in the follow-up experiment were also randomly assigned to either saline (N = 10) or IN-OT treatment (N = 8) prior to the retest. Immediately prior to the 5-10 min habituation preceding the retest, females were given either IN-OT or control saline, while all males were administered saline, as this time course has previously been shown to result in behavioral effects in California mice [40]. To administer, a mouse was scruffed and 25 μl of solution was administered to the nostrils using a blunt needle attached to cannula tubing at the end of a Hamilton syringe. Individual droplets were beaded at the end of the syringe, applied to the surface of the nose, and allowed to absorb into the nasal mucosa. Administration lasted less than 30 s per animal. Only females were given IN-OT because of unpublished data suggesting that IN-OT shifts the convergence/division of labor ratio for pairs defending their territories when given to females, but not males [55].

### USV analysis

We recorded USVs with two ultrasonic microphones (Emkay/Knowles FG, detection range 10-120 kHz) placed 85 cm apart at opposite corners of the apparatus and 30 cm from the apparatus floor, with one microphone placed in the approach chamber and the other placed in the avoid chamber. Microphone channels were calibrated to equal gain (−60 dB noise floor). WAV files were produced using RECORDER software (Avisoft Bioacoustics, Berlin, Germany). Recordings were made using a 250 kHz sampling rate with 16-bit resolution and spectrograms were produced with a 512 fast Fourier transform made using Avisoft SASLab Pro (Avisoft bioacoustics). USVs were differentiated by visual and auditory inspection of WAV files with sampling rates reduced to 4% of normal speed (11025 kHz). Beyond barks, California mice produce a variety of ultrasonic vocalizations (USVs) [39] important for communication and behavioral coordination [56]. Because barks are only typically produced when aggressive physical contact is made [16] and were not produced by mice in this study, two call types, sustained vocalizations (SVs) and sweeps, were analyzed, because they may play a role in pair bond formation and maintenance [48,57,58]. SVs are low-bandwidth calls with low modulation, a peak frequency of 20 kHz and a duration of 100 to 500 ms for each individual syllable (Fig. 1B). Shorter SVs are used during aggressive encounters [16]. Sweeps are of relatively short durations of 30 to 100 ms with upwards and downwards modulation ranging from 25 to 100 kHz and potentially including multiple inflection points [16] (Fig. 1C). Because audio was recorded for both pair members simultaneously, it was impossible to differentiate calls by each individual so vocalizations were analyzed at the pair level. Both the total number of USV calls produced and the proportion of each USV individual call type produced relative to all call type production were analyzed within this dataset. Four audio files were unusable due to recording setup error (mismatched saline pairs: 11; matched saline pairs: 8; mismatched OT pairs: 14; matched OT pairs: 9). The length of SVs was also analyzed, and eight pairs that did not produce SVs were removed from this analysis only (mismatched saline pairs: 11; matched saline pairs: 5; mismatched OT pairs: 12; matched OT pairs: 6).

### Statistics

All statistics were analyzed using R (version 3.6.2). We analyzed the female-male approach difference before pairing by taking the absolute value of the difference between female (F) and male (M) approach, |F_1_-M_1_| and regressing it on pair type (mismatched or matched) in a general linear model (GLM). We analyzed the degree to which pairs converged by taking the initial difference between pair members, |F_1_-M_1_|, and subtracting it from the difference at the second test, |F_2_-M_2_|, and regressed it on pair type, treatment type (saline or IN-OT), and their interaction in a GLM. We analyzed individual changes by subtracting initial approach from approach at the second test (F_2_-F_1_ and M_2_-M_1_) and regressed it on pair type, treatment type, and their interaction in a GLM. We likewise analyzed the amount of time pair members spent within the same chamber using pair type, treatment type, and their interaction as factors in a GLM. For changes in vocalizations, we summed vocalizations produced by pairs before pairing and subtracted them from the vocalizations produced in the second test, again using pair type, treatment type, and their interaction as factors in a GLM. For the proportion of sustained vocalizations (SVs), we looked only at vocalizations produced at the post-pairing test, because there were so few vocalizations produced prior to pairing. We analyzed the proportion of SVs using a GLM with pair type, treatment, and their interaction as factors. We used linear regressions to test if behavior predicted call type proportion. For the separate experiment using unpaired females, we regressed second approach on initial approach, and also analyzed the effects of treatment on approach during the second test, using a GLM. P-values were Holm-Bonferroni corrected for multiple comparisons where appropriate.

## Results

### Mismatched pairs had a greater difference in approach scores before pairing than did matched pairs

Prior to pairing, a wide range in approach to bark playbacks was found in females (33.43 ± 4.07 s s.e.m.) and males (37.96 ± 4.19 s s.e.m.), with a general linear model (GLM) of approach regressed on sex revealing no mean difference between sexes (F(1,90) = 0.597, p = 0.442; Fig. 2B). Males and females were paired based on their approach, whereby “mismatched” pairs had ≥ 30 s difference in approach score while matched pairs had ≤ 10 s in their approach score (“mismatched”: N=16 pairs female lower than male, N=13 pairs male lower than female; “matched”: N=8 pairs female lower than male, N=9 pairs male lower than female; Fig. 2C). As expected, a GLM revealed a statistically significant gap between the approach difference for females and males of mismatched (50.59 ± 3.11 s s.e.m.) and matched (5.71 ± 0.83 s s.e.m.) pairs (F(1,44) = 118.100, p < 0.001; Fig. 2D).

### IN-OT drove mismatched pairs to become more similar than saline mismatched pairs

To determine how IN-OT affected pair convergence, we first assessed the change in approach score difference for matched and mismatched pair mates from the before-pairing test to the after-pairing test. A GLM revealed a significant interaction between pair type (matched or mismatched) and treatment (saline or IN-OT) (F(1,42) = 4.189, p = 0.047; Fig. 3A; Table 1A). Controlling for treatment effects, mismatched pairs decreased their approach score difference more than matched pairs from before-pairing to after-pairing, indicating that mismatched pairs become more similar, while matched pairs remain similar (F(1,42) = 58.346, p < 0.001). Relative to saline, IN-OT treatment resulted in a further decrease in approach score difference for mismatched but not matched pairs, as shown by a simple effect of treatment within mismatched pairs, suggesting that IN-OT drove mismatched pairs to become more similar (F(1,42) = 11.539, p = 0.002).

**Table 1.**
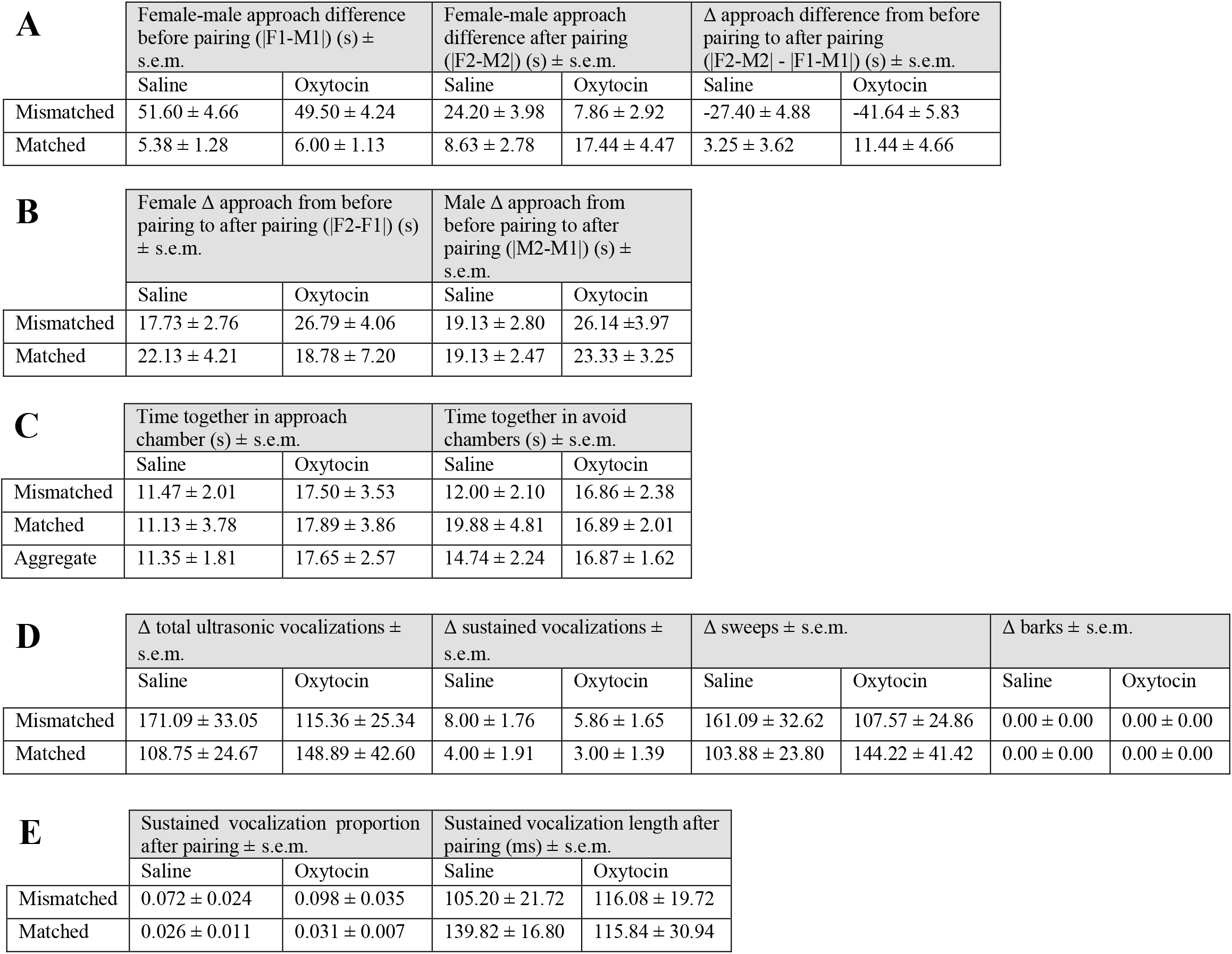
Descriptive table of mean group values. **A**. Approach differences between paired mates. **B**. Approach differences for individuals. **C**. Time spent together by pairs. **D**. Changes in aggregate vocalizations from before pairing to after pairing. **E**. Proportion of ultrasonic vocalizations that were sustained vocalizations after pairing.

**Figure 3.**
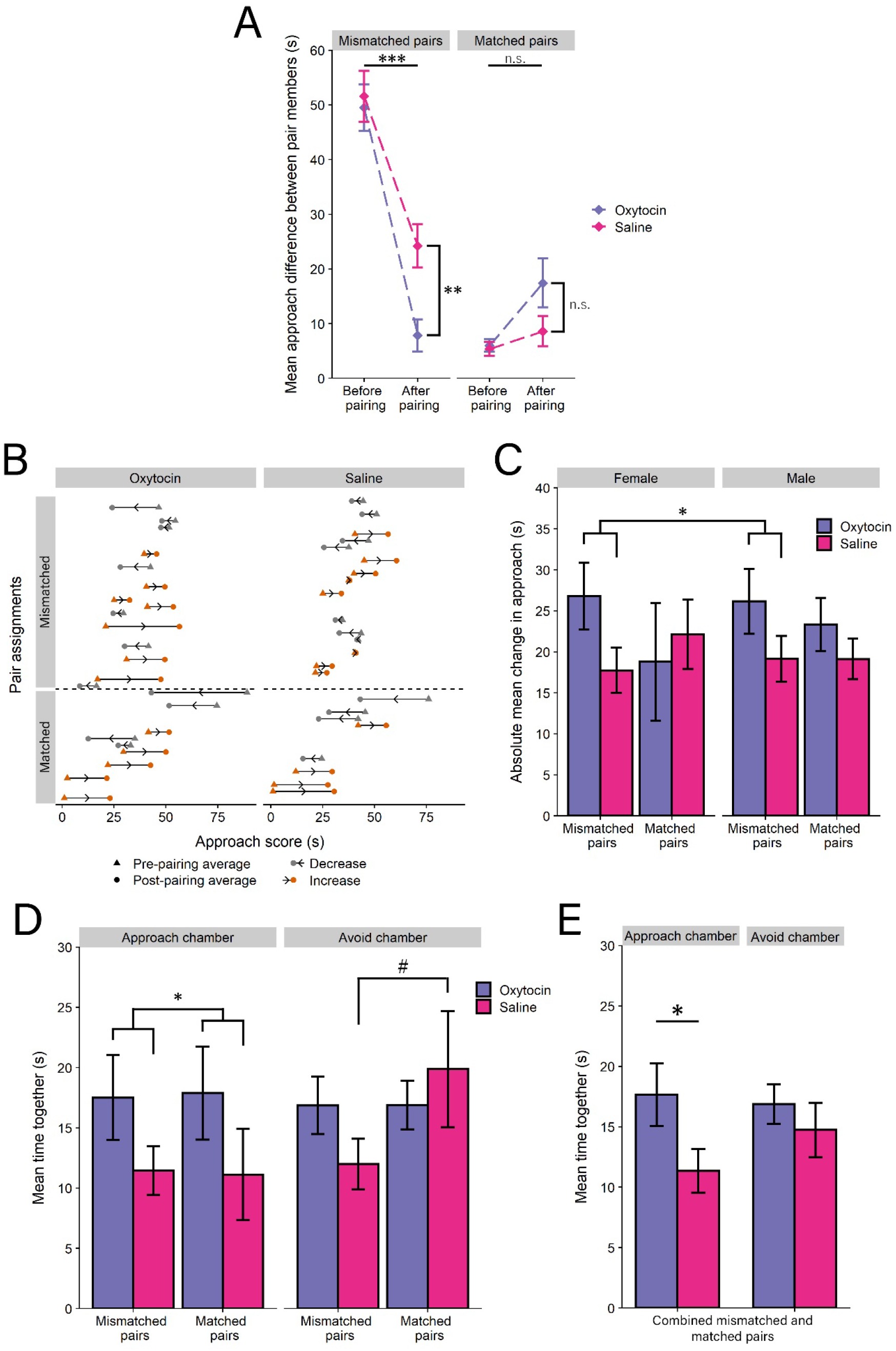
IN-OT drove mismatched pair mates to converge more in their social approach and pairs to increase their joint approach. **A**. Mismatched pair mates became more matched, an effect enhanced by OT, while matched pairs remained matched and were unaffected by OT. **B**. Within-pair average approach increased and decreased by pair for mismatched and matched OT and saline groups. **C**. Females and males of OT mismatched pairs changed their approach more than females and males of saline mismatched pairs. **D**. IN-OT increased the amount of time OT pair mates spent together in the approach chamber. **D**. Time together in the approach chamber and avoid chamber, aggregated for all pairs. (n.s. = not significant; # = p < 0.10; * = p < 0.05; ** = p < 0.01; *** = p < 0.001). Error bars are ± s.e.m.

From pre-pairing to post-pairing, mismatched and matched IN-OT and saline groups all contained pairs that had pair-averaged approach scores that decreased and pair-averaged approach scores that increased, suggesting that convergence may be driven differently by individuals within pairs (Fig. 3B). To determine whether males and females contribute differently to the overall convergence of pair approach scores, we regressed change in approach from the before-pairing test to the after-pairing test for each individual on the interaction of pair type, treatment, and sex. This GLM revealed no 3-way interaction (F(1,84) = 0.693, p = 0.408; Fig. 3C), however, IN-OT individuals in mismatched pairs had a greater change in approach from the before-pairing test to the after-pairing test than did saline individuals in mismatched pairs when controlling for sex, suggesting that the convergence in IN-OT mismatched pairs emerged from changes in approach by both sexes (F(1,84) = 5.257, p = 0.0244; Table 1B).

### IN-OT increased the amount of time pairs spent in joint approach

Complementing the analyses on the degree to which pair mates changed their behavior, we also measured time spent together as an indicator of coordination. Analysis of the amount of time paired mates spent together in either the approach chamber (the chamber closest to the bark playback speaker) or the avoid chamber (the chamber closest to the ambient noise speaker), via a 3-way interaction of pair type, treatment, and location, revealed that IN-OT spent significantly more time together in the approach chamber than saline pairs (F(1,84) = 4.311, p = 0.041) but not in the avoid chamber (F(1,84) = 0.092, p = 0.762), and no main effect of location (F(1,84) = 0.768, p = 0.383), pair type (F(1,84) = 1.645, p = 0.203), or the full interaction (F(1,84) = 0.967, p = 0.328; Fig. 3D; Table 1C). Additionally, there was a nonsignificant effect of pair type in the avoid chamber for saline groups (F(1,84) = 3.016, p = 0.086). A separate analysis that combined mismatched and matched pairs into a single category also revealed a significant main effect of IN-OT (F(1,88) = 4.066, p = 0.047), with a significant simple effect whereby IN-OT increased the amount of time pairs spent together in the approach chamber (F(1,88) = 4.543, p = 0.036) but not the avoid chamber (F(1,88) = 0.519, p = 0.473; Fig. 3E). There was also no effect of location (F(1,88) = 0.389, p = 0.535) or the interaction of treatment and location (F(1,88) = 0.996, p = 0.321).

### Mismatched pairs produced a higher proportion of sustained vocalizations after pairing, which was not affected by IN-OT

To determine how pair type and IN-OT affected vocal production, we assessed the number and type of USVs produced in the pre-pairing and post-pairing tests. A GLM revealed a significant increase in the total number of vocalizations produced by pairs relative to the sum of vocalizations for pair members before pairing (F(1,38) = 70.267, p < 0.001), but no significant interaction between pair type and treatment (F(1,38) = 2.182, p = 0.148; Fig. 4A). Similarly, a GLM for number of sweeps (F(1,38) = 66.182, p < 0.001) indicated significant increases in sweep production after pairing, but no significant interaction (F(1,38) = 2.184, p = 0.148; Fig. 4B). A GLM for number of sustained vocalizations (SVs) revealed a significant increase in SV production after pairing (F(1,38) = 35.168, p < 0.001) as well as a nonsignificant increase in SV production by mismatched pairs (F(1,38) = 3.801, p = 0.059), but no main effect of treatment (F(1,38) = 0.799, p = 0.377) or significant interaction (F(1,38) = 0.106, p = 0.747; Fig. 4C; Table 1D). Taken together, this suggests that pairing increases the total number of sweeps and SVs that are produced, and that pair type and IN-OT do not affect the total number of either sweeps or SVs.

**Figure 4.**
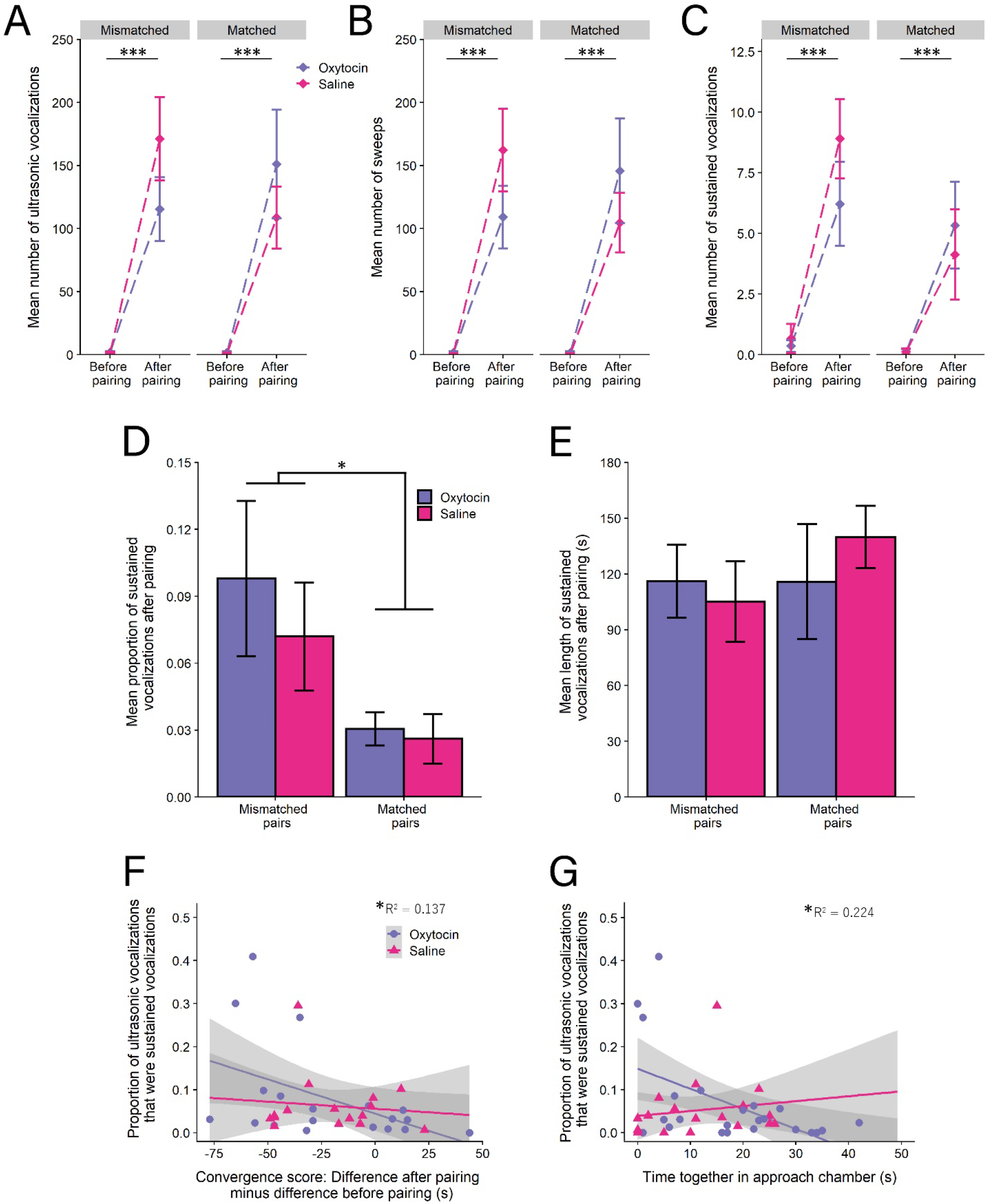
Pair type and IN-OT altered the production of ultrasonic vocalizations. **A**, **B**, **C**. Pairing resulted in a profound increase in the number of ultrasonic vocalizations produced, including sweeps and sustained vocalizations, across pair type and treatment. **D**. Mismatched pairs produced a greater proportion of sustained vocalizations. **E**. Length of sustained vocalizations did not differ by pair type or treatment condition. **F**.The degree to which pair mates converged predicted the proportion of sustained vocalzations produced after pairing. A negative number on the x-axis shows pair mates decreased the difference in approach score from the first test to the second test, indicating convergence. **G**. IN-OT and time spent together in the approach chamber negatively predicted the proportion of SVs produced. (* = p < 0.05; *** p < 0.001). Error bars are ± s.e.m.

We next assessed whether other features of SV call production were affected by pair type or IN-OT. A GLM revealed that mismatched pairs significantly increased the proportion of SVs produced after pairing (as used in [57]) (F(1,38) = 4.141, p = 0.048), but that there was no significant interaction with IN-OT (F(1,38) = 0.298, p = 0.588; Fig. 4D, Table 1E). A GLM indicated no significant effect of pair type (F(1,31) = 0.475, p = 0.496), treatment (F(1,31) = 0.069, p = 0.795), or interaction (F(1,31) = 0.489, p = 0.490) on SV length (Fig. 4E, Table 1E). These results indicate that SV proportion may be involved in pair convergence.

### Sustained vocalization proportion positively correlated with pair similarity and negatively correlated with time IN-OT-treated pairs spent together in the approach chamber

To determine whether SV proportion after pairing was associated with pair coordination, we correlated SV proportion of pairs with convergence and treatment, and then joint approach and treatment. A GLM revealed a significant main effect of the degree of convergence (|F2-M2| − |F1-M1|) on the proportion of SVs produced after pairing (F(1,38) = 4.476, p = 0.041; Fig. 4F), showing that the degree to which pairs become more similar is positively correlated with SV production. However, there was no significant interaction between pair type and treatment (F(1,38) = 0.595, p = 0.445) or main effect of treatment (F(1,38) = 0.105, p = 0.748), suggesting that IN-OT did not have an effect on the relationship between SV proportion and convergence.

A separate GLM revealed a significant interaction between the amount of time pairs spent together in the approach chamber and treatment type on the proportion of SVs produced (F(1,38) = 5.000, p = 0.031, Fig. 4G), with a significant simple effect of IN-OT (F(1,38) = 10.440, p = 0.003) but not saline (F(1,38) = 0.280, p = 0.600), indicating that increased time spent together in the approach chamber for IN-OT pairs was associated with decreasing SV proportion.

### Unpaired females respond to bark playbacks in a consistent manner

In order to demonstrate that approach behavior is repeatable and to investigate the effects of the IN-OT treatment in unpaired individuals, a separate cohort of unpaired females was tested for initial response and response 13-17 days later, the same length of time as that between the pre-pairing and post-pairing test for paired mice. A GLM revealed that initial approach reliably predicts later approach (F(1,14) = 14.59, p = 0.002; Fig. 5A), with no significant main effect of treatment (F(1,14) = 0.313, p = 0.585) or interaction (F(1,14) = 0.227, p = 0.641). A separate GLM indicated no effect of treatment on approach during the second test (F(1,16) = 0.415, p = 0.529; Fig. 5B).

**Figure 5.**
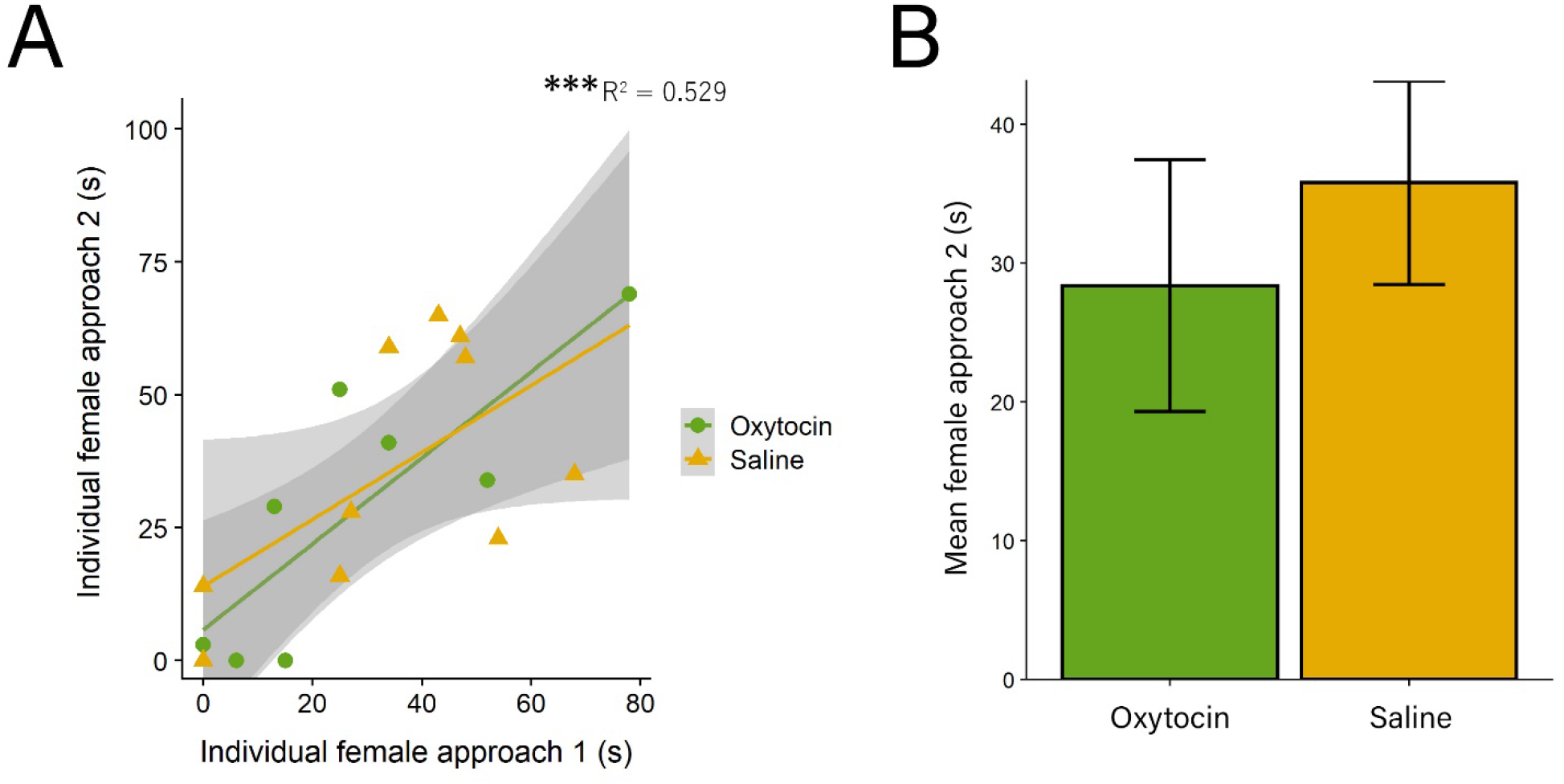
A separate cohort of unpaired females showed high test-retest reliability. **A**. Unpaired females showed reliable response to the playback approach test 13-17 later. **B**. IN-OT did not significantly change female approach at the second test. (*** = p < 0.001). Error bars are ± s.e.m.

## Discussion

The social salience hypothesis argues that oxytocin (OT) attunes individuals to social context [24], possibly explaining why OT is able to induce both social avoidance [19] and social approach [41]. However, the roles of affiliative bonds in this process have remained elusive. Here we used a pre-post pairing design and found that OT facilitates pair-bonding-induced changes in social behavior and coordination. Intranasal OT (IN-OT), when given to female California mice, produced pair-level shifts in social approach of an aggressive bark stimulus driving mismatched pair mates to become more similar. Analysis of time spent in proximity showed that IN-OT drove pairs to spend more time together approaching the stimulus, likely producing the shift towards pair similarity and indicating coordinating effects of OT beyond bonding in these already bonded pair members. Recordings of USVs revealed that IN-OT drove the negative correlation between the ratio of SVs and time spent together in approach. Finally, unpaired females demonstrated a high degree of repeatability in their approach, which was not affected by IN-OT. These data suggest that the social context of pair bonding alters social approach, that the degree to which an individual alters their social approach depends on their partner, and that OT facilitates pair-bonding-dependent changes in social approach. Our results are consistent with a social convergence theory, whereby OT unites pair-bonded individuals in the face of challenge, and that this coordination may be related to changes in the proportions of vocalization types produced.

Recently, two other Peromyscus species were shown to have repeatability in behavior and correlation between boldness behaviors [59], and other consistent individual differences have been found in Peromyscus species [60], including California mice [39]. We extend these findings to consider the change in behavior type resulting from the presence of a pair member, and vocalizations and OT signaling as behavioral and hormonal mechanisms, respectively, under affiliative social contexts. We observed that IN-OT given to females increased convergence in approach scores for initially mismatched pairs but had no effect on either the convergence of pairs that were initially matched or on total time unpaired females spent approaching the simulated intruder, suggesting that OT changes in behavior may be related to a motivation to maintain partner proximity in response to a single stimulus. While convergence has been shown to be asymmetrical in convict cichlids, with the more reactive partners being more flexible than proactive partners [43], we show that in California mice both individuals with low and high initial levels of social approach change their behavior in response to pairing. Our results can be interpreted as adjustments by both pair members to become more similar in behavioral type to their partners, and not solely within-individual variation in approach response from the pre-pairing to the post-pairing test, because unpaired individuals showed repeatable responses to the vocalization playback stimuli. Taken together, our study provides experimental evidence of pair type-dependent changes to social behavior that are facilitated by female OT signaling, and is consistent with prior evidence of convergence by mismatched pairs [38, in review]. Moreover, it suggests that although individual unpaired females demonstrate reliable and consistent approach behavior even in response to IN-OT, the change in approach behavior induced by pairing mismatched individuals is malleable in response to IN-OT.

Several hypotheses emerge for why IN-OT drives changes in behavior. First, it is possible that social approach reflects underlying differences in stress and stress susceptibility, as the stressed state of an individual influences both social approach and OT signaling. Social defeat stress decreases OT receptor (OTR) gene expression in female California mice [19] and reduced OTR binding in the NAc corresponds to reduced social approach following defeat [61]. Moreover, plasma OT levels have been positively linked to attachment anxiety in romantic human relationships [62], and pair bonding influences affective state across taxa [63,64]. Because we did not directly measure stress or anxiety, we are unable to correlate individual differences in stress response to social approach or pair bonding-induced convergence. However, while IN-OT may have influenced anxiety-like behavior, the lack of IN-OT to change social approach in unpaired individuals suggests no baseline interaction between stress and OT.

Another hypothesis for why IN-OT drove changes in behavior is that it increased the motivation to bond. While it is possible that the behavioral convergence reflects a motivation to stay closer together to maintain the pair bond, the second test occurred after the pairs were stably bonded. IN-OT likely induced coordination of behavior beyond bonding, as evidenced by the increase in time IN-OT pairs spent jointly approaching the stimulus, but not jointly avoiding. Further evidence is suggested by the vocalization data: In this study we tracked two call types, sustained vocalizations (SVs) and sweeps, because they may play a role in pair bond formation and maintenance [58]. While mismatched pairs produced a higher proportion of sustained vocalizations, indicating a potential behavioral mechanism underlying this coordination, IN-OT drove a negative correlation between SV proportion and time spent together in the approach chamber, suggesting that OT may be involved in the relationship between decreased SV production and behavioral convergence. We found no difference in SV length between groups, which has previously been shown to be a relevant characteristic for bonding pairs, with longer SVs produced over the first week of pairing [56]. However, different results might have occurred if we had recorded individual USVs as opposed to pair USVs. Moreover, the SVs in this study were relatively short compared to those found by other studies (an average of about 115 ms compared to, for example, an average of about 175 ms in [48]), but similar to those observed during aggressive encounters (for example, an average of about 105 ms in [16]) which may indicate negotiation or negative affect by pairs in this novel context.

Finally, another hypothesis for why IN-OT drove changes in behavior relates to pair coordination and synchrony. Synchrony, an emergent behavior that occurs between two or more individuals, such as eye contact, singing, or movement, is linked to greater affiliation and cooperation [65]. California mice are socially monogamous and form strong affiliative bonds that last for life [66]; OT may increase the ability of pair-bonded individuals in monogamous, biparental species to perceive a partner’s intent and thus provide complimentary behaviors to efficiently address environmental challenges such as foraging, defending against predators, and taking care of young.

It is interesting to note that IN-OT did not increase sweeps as it has previously been shown to do so when given to mothers interacting with their pups [40]. This effect was context-dependent such that OT increased sweeps in mothers only when mothers were in physical contact with their pups. In the current study we observed an increase in sweeps in response to pairing as expected by [48], but IN-OT did not increase production of sweeps. The reason for this is unknown, however, it is possibly because the playbacks resulted in a less affiliative interaction, or that IN-OT effects are restricted to mother-pup interactions.

Overall our results suggest that OT modulates social approach in a social context-dependent fashion by driving initially mismatched pairs to become more matched by increasing time spent in joint navigation, consistent with previous findings that OT signaling is critical to both bonding [67] and social coordination [23]. Taken together these results suggest that affiliative social context is critical for OT-regulated social approach and vigilance. An important direction for future studies is determine the neural circuitry overlap between social approach and affiliative bonding. Our results are in line with calls for researchers to consider social context when studying behavioral types [68,69].

## Acknowledgements

Research was conducted at the University of Wisconsin-Madison, which occupies the ancestral Ho-Chunk land known as Teejop. Following an 1832 treaty, federal and state governments repeatedly, but unsuccessfully, sought to forcibly remove the Ho-Chunk from Wisconsin [70]. As members of a land grant institution we directly benefit from land theft, and we challenge ourselves and others to reflect on the perpetuation of the colonialist roots of western scientific progress. A. Auger, L. Riters, C. Guoynes, and C. Malone provided manuscript feedback, and Z. Herro and T. Nguyen helped with data collection. We also thank the UW-Madison animal research technicians. Research was supported by the National Science Foundation (IOS-1946613 and DGE-1747503).

## Contributions

PKM, NSR, and CAM designed the study, PKM and JS conducted the experiments and data collection, PKM analyzed data, PKM wrote the initial draft of the manuscript, CAM and NSR provided guidance during analyses, and all authors contributed to the writing and review of the manuscript.

## Data accessibility

Data available from the Open Science Framework: https://osf.io/2a8gm/

